# The boundaries of State-Space Granger Causality Analysis applied to BOLD simulated data: a comparative modelling and simulation approach

**DOI:** 10.1101/2020.04.10.033258

**Authors:** Tiago Timóteo Fernandes, Bruno Direito, Alexandre Sayal, João Pereira, Alexandre Andrade, Miguel Castelo-Branco

## Abstract

**Background:** The analysis of connectivity has become a fundamental tool in human neuroscience. Granger Causality Mapping is a data-driven method that uses Granger Causality (GC) to assess the existence and direction of influence between signals, based on temporal precedence of information. More recently, a theory of Granger causality has been developed for state-space (SS-GC) processes, but little is known about its statistical validation and application on functional magnetic resonance imaging (fMRI) data.

**New Method:** We implemented a new heuristic, focusing on the application of SS-GC with a distinct statistical validation technique - Time Reversed Testing - to generative synthetic models and compare it to classical multivariate computational frameworks. We also test a range of experimental parameters, including block structure, sampling frequency, noise and system mean pairwise correlation, using a statistical framework of binary classification.

**Results:** We found that SS-GC with time reversed testing outperforms other frameworks. The results validate the application of SS-GC to generative models. When estimating reliable causal relations, SS-GC returns promising results, especially when considering synthetic data with an high impact of noise and sampling rate.

**Conclusions:** SS-GC with time reversed testing offers a possible framework for future analysis of fMRI data in the context of data-driven causality analysis.

**Highlights:**

- State-Space GC was combined with a statistical validation step, using a Time Reversed Testing.
- This novel heuristic overpowers classical GC, when applied to generative models.
- Correctly identified connections between variables increase with the increase of number of blocks and number of points per block.
- SNR and subsampling have a significant impact on the results.

## 1. Introduction

The characterization of the connectivity structure between brain regions, in health and disease, is a matter of considerable neuroscientific interest. Connectivity measures provide information about the interaction between brain regions and help to describe brain structure and function; these are believed to represent potentially reliable biomarkers of neuronal dysfunction (Fox, 2010; Pievani et al., 2014).

Several methods have been proposed to assess connectivity from neural time series and infer the underlying neural associations among segregated brain regions (Stephan and Roebroeck, 2012; Valdes-Sosa et al., 2011). These methods can be divided into functional connectivity (FC) (directed functional connectivity (dFC) and undirected) and effective connectivity (EC).

FC is defined in terms of statistical dependencies between segregated neuronal regions (dFC appeals to the temporal precedence, i.e. reflects an underlying dynamical process, in which causes precede consequences), whereas EC refers to the influence that one neural system exerts over another. See (Friston et al., 2013) for a full comprehensive description of these concepts.

dFC and EC have been widely studied in neurophysiological data from different imaging modalities such as functional magnetic resonance imaging (fMRI) (Ghasemi and Mahloojifar, 2013; Kadosh et al., 2016; Krueger et al., 2011; Schippers et al., 2011), electroencephalography (EEG) (Barrett et al., 2012; Flamm et al., 2013) and Local Field Potentials (LFP) (Brovelli et al., 2004; Gaillard et al., 2009). Two of the most common connectivity measures are Granger Causality (GC) (Granger, 1969), and Dynamic Causal Modelling (DCM) (Friston K.J., Harrison L., 2003).

Granger Causality Mapping (GCM) (Roebroeck et al., 2005), a measure of dFC, is a data-driven method that uses GC to assess the existence and direction of influence between signals, based on the temporal precedence of information. Its application to fMRI has been object of intense debate within the scientific community (David et al., 2008; Seth et al., 2015; Valdes-Sosa et al., 2011; Wen et al., 2013), raising a number of technical issues (Mill et al., 2017; Seth et al., 2013). These authors investigated the influence of different model parameters on the computation of GC.

The Hemodynamic Response Function (HRF) (i.e. the function that relates the fMRI signal to neural activity) represents a source of delay and variability between nodes/regions (Valdes-Sosa et al., 2011) that may compromise the estimation of GC. Seth and colleagues (Seth et al., 2013) addressed this issue, reinforcing that the causal influences measured on Blood Oxygenated Level-Dependent (BOLD) signal are invariant to the HRF for a very high sampling rate, but influenced by severe downsampling, as is typical for current fMRI data.

Other parameters have also been investigated, including repetition time (TR) i.e. the time required to sample two consecutive measures, in the order of the second, signal to noise ratio (SNRs) and neuronal delay (Mill et al., 2017; Rodrigues and Andrade, 2014). The results support that, if certain conditions are met, GC is applicable in the context of fMRI.

The underlying neural processes commonly studied with fMRI vary in the scale of the millisecond and are sampled in the scale of the second. This measure of neuronal activity roughly represents a low pass filter of LFPs (Logothetis et al., 2001), showing that neural patterns can also be found in the BOLD signal.

The subsampling associated with the fMRI data raises questions regarding the possible distortions (e.g. preservation or reversal) on the GC directionality (Barnett et al., 2018; Solo, 2016). (Seth et al., 2013) suggested that sampling might affect the conclusion drawn on directionality. There are some indications of successful estimation of causal connectivity; (Kadosh et al., 2016; Wen et al., 2013) outlined causal influences among regions, arguing that even considering low sampling, the results were physiologically significant.

The most common approach to determine GC is to consider a linear autoregressive (AR) model. However, data frequently contain a Moving Average (MA) component, either intrinsic to the underlying signal or induced by data filtering, acquisition, sampling and preprocessing. These components make data ill-suited for auto-regressive (AR) modelling (Barnett and Seth, 2015).

To overcome these limitations, (Barnett and Seth, 2015; Solo, 2015) proposed a novel computational approach, State-Space GC (SS-GC), tackling the issues of measurement noise and downsampling in fMRI data. The method addresses the assumptions of linearity, stationarity and homoscedasticity, and requires alternative techniques to permutation or blockwise randomization to evaluate statistical significance.

Winkler and Haufe (Haufe et al., 2013, 2012; Winkler et al., 2016) proposed a statistical framework based on a time reversed time-series. Reversed time series share weak asymmetries (spectral densities, cross-covariance) with the original data but reduce strong asymmetries (temporal ordering, that leads to causal relations), alleviating the problem of spurious causality associated with other approaches.

In this paper, we analyzed connectivity methods on synthetic data with a well defined architecture of influence between signals. We considered temporal discontinuities, representative of block-based and task design paradigms. Firstly, we aimed to define the optimal combination of GC estimation method and statistical framework, exploring two synthetic datasets.

We hypothesized that the combination of SS-GC with a statistical validation based on time reversed testing would represent a methodological alternative.

Based on the optimal combination, we explored a wider range of parameters (such as number of data points, number of blocks, sampling frequency, SNR, number of network nodes and mean correlation, i.e. mean pairwise correlation between variables), and discussed scenarios of possible application to fMRI data.

## 2. Materials and Methods

In this section we summarize Granger causality and present an overview of the statistical framework.

### 2.1. Granger-Causality as a time-lagged measure

GC can be operationalized considering a vector autoregressive (VAR) model (Barnett and Seth, 2014).

Considering a n-dimensional multivariate time series *u*_*t*_, where *t* represents each time point, a *p*^*th*^-order model is defined as:

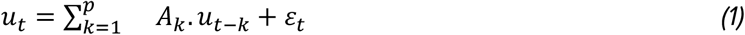

where *A* represents the regression coefficients and *ɛ* the residual matrix, which represents a *n*-dimensional white noise process.

There are several approaches to determine the appropriate model order which is pivotal to optimize model fit. To this end, information theory concepts such as the Akaike Information Criterion or the Bayesian Information Criterion can be used.

GC is a method that can be used to identify directed functional connectivity between time series. A time series Y is said to G-cause a time series X if the past of Y contains information that improves the prediction of X over the use of all available information apart from Y.

Let us consider a three-dimensional model, X, Y and Z. The causality from Y to X, in this case, is conditioned by a third variable, Z. The causality *Y* → *X*|*Z*, can be defined as the logarithm of the covariance matrix of the full and of the partial system, as follows:

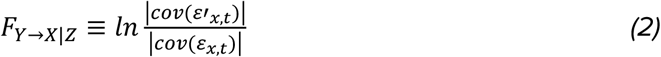

where *ɛ*_*x,t*_ represents the residuals of the VAR model considering the three-dimensional model and *ɛ*′_*x,t*_ represents the residuals of the two-dimensional model (with only *X* and *Y*), by doing so one “conditioning out” the common dependencies. This is the approach used in the Multivariate Granger Causality Toolbox - MVGC (MATLAB implementation described in (Barnett and Seth, 2014))

### 2.2. State-space models

Let us consider a system composed on two discrete-time, real-valued vectors *z*_*t*_ = [*x*_*t*_, *y*_*t*_] , −∞ < *t* < +∞.

The SS is a mathematical formulation of a system describing the relation between a set of input, output and state variables. Equations 3 and 4 represent a general constant parameter SS model:

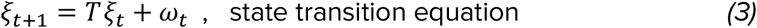

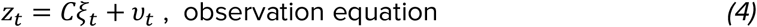

where *ξ* is a *n*-dimensional state variable (unobserved), *ω*_*t*_, *v*_*t*_ zero-mean white noise processes, *T* the state transition matrix and *C* the observation matrix. The model can be referred to as (*T*, *C*, [*QRS*]), where Q, R and S are based on 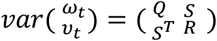.

The Kalman filter theory applied to SS formulation, provides small corrections to the model predictions, that allows us to define the previous model as:

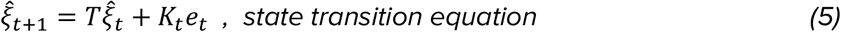

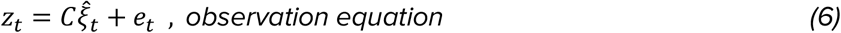

where 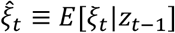, *K* is the Kalman gain, and *e*_*t*_, also known as the innovations, are defined as 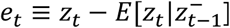 and constitute a white noise process (for additional details, Barnett and Seth, 2015; Solo, 2015; Bishop, 2006). The time-domain GC from Y to X conditional to Z would then be:

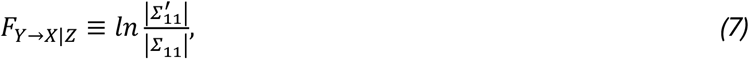

where 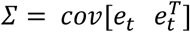 and 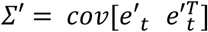, full and of the partial system, respectively.

### 2.3. Statistical inference

Considering the VAR model, the GC is estimated considering a test statistic for the null hypothesis of no interaction (zero GC)

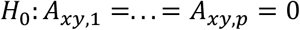

where *A*_*xy*_ represents the VAR coefficients matrix for the contribution of the past of *Y* on the present of *X*. Alternatively, *Y* G-causes *X* if the full model provides a significantly better model than the reduced model, at a given significance level. Since the reduced model is ‘nested’ in the full model, the appropriate test for *H*_0_ is the likelihood ratio (under normality assumption of the residuals *ɛ*′_*x,t*_ and *ɛ*_*x,t*_) *F*_*Y*→*X*|*Z*_. Moreover, the estimates are asymptotically equivalent to those obtained by ordinary least squares algorithm which represents an easy computational approach (Barnett and Seth, 2014).

Statistical inference for SS models presents different challenges, since the assumptions for VAR models are not valid (Barnett and Seth, 2015).

A possible approach for statistical validation is based on surrogate data techniques (creation of time series with disrupted causal relationships). After constructing an ensemble of independent surrogates, we determine an empirical distribution for the null hypothesis. If the value estimated for the original time series is significantly higher than the empirical distribution of values obtained for the surrogates, the corresponding underlying null hypothesis can be rejected.

The creation of permutation surrogates (PS) is based on the random shuffling of segments of the original time series. In order to maintain the causal information, the data must be divided into equally-sized segments, larger than the model order (Anderson and Robinson, 2001).

Time reversed testing is an alternative approach based on time reversal of the original time-series which keeps weak asymmetries, including features such as spectral densities and cross-covariance, but reduces strong asymmetries (e.g. temporal ordering) that lead to causal relations. This approach has been proposed by (Haufe et al., 2012) to avoid false detection of causal interactions, while assessing GC applied to the original time series and time reversed signals.

### 2.4. Synthetic data

The synthetic data generated explores several parameters in order to identify an optimal framework for GC computation, ultimately discussing its applicability to the fMRI data scenario. The explored parameters were the number of samples (time series length), data partition in blocks, downsampling (sampling frequency of the signal), signal to noise (SNR), number of variables (or network nodes) and their mean correlation.

#### 2.4.1. Generative models

The synthetic datasets were based on multivariate auto-regressive (MVAR) models with different numbers of variables, considering different patterns of pairwise correlation and auto-correlation.

The data were generated with a sampling frequency of 1000 Hz. Taking into account that the neural processes occur in the time-scale of milliseconds (Schippers et al., 2011), we considered a neural delay *l* of 20 ms.

Model A was based on an MVAR model with 5 variables (Baccalá and Sameshima, 2001). This MVAR model, which contains mutual influences between variables, unidirectional and bidirectional connections, is illustrated in Figure 1 (see supplementary equation S.1 for additional details).

**Figure 1.**
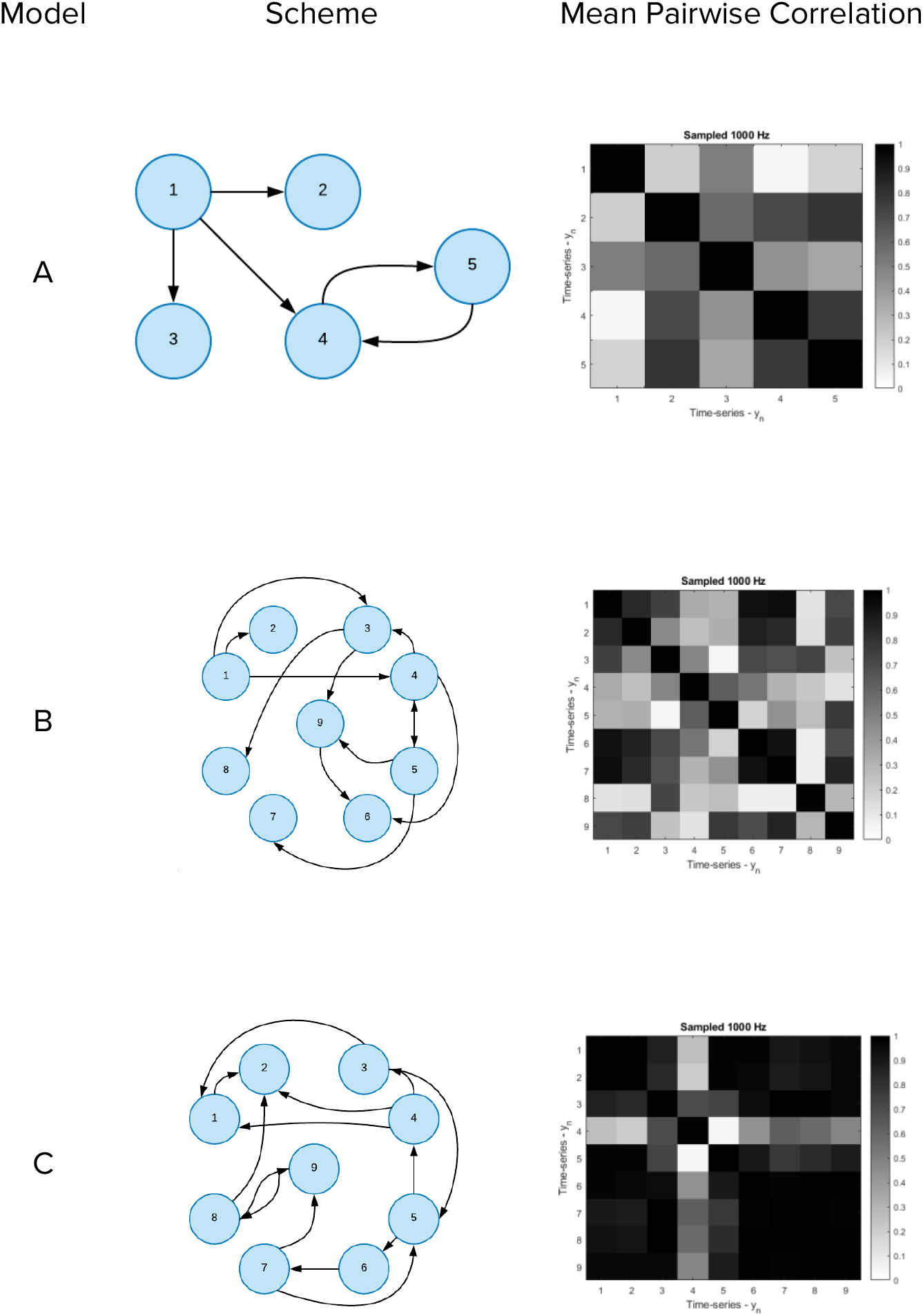
Model A, B and C, conceptual representation and mean pairwise correlation matrix; A - 5 MVAR model with a mean value of 0.0854; B - 9 MVAR model with weak mean pairwise correlation c with mean value of 0.2165; C - 9 MVAR model with strong mean pairwise correlation with mean value of 0.8676;

Model B was synthesized based on an MVAR model with 9 variables (Figure 1.B), presenting weak mean pairwise correlation among nodes (see supplementary equation S.2 for additional details). Model C (shown in Figure 1.C) was based on the same model but presented strong mean pairwise correlation between variables - the rationale here was to compare the behavior of the connectivity estimation framework considering MVAR models with different mean pairwise correlation values (see supplementary equation S.3 for additional details). In both models there are unidirectional connections and bidirectional connections

#### 2.4.2. Variable HRF convolution

The output of the generative models was convolved with an Hemodynamic Response Function (HRF). We used the default HRF implemented in SPM8 (Friston et al., 2007) to generate two sets of curves (one for each number of model variables - Figure 2). HRF times-to-peak are deliberately confounded with the underlying neural delays of all models, meaning that the nodes which drive flow of information could actually have their HRF peaks after the HRFs of their targets, which allows for a conservative approach. The first 50 seconds of each dataset were excluded to remove the HRF ‘burn-in’ effect (Seth et al., 2015, 2013). The outcome generative models can be depicted on Figure 3.

**Figure 2.**
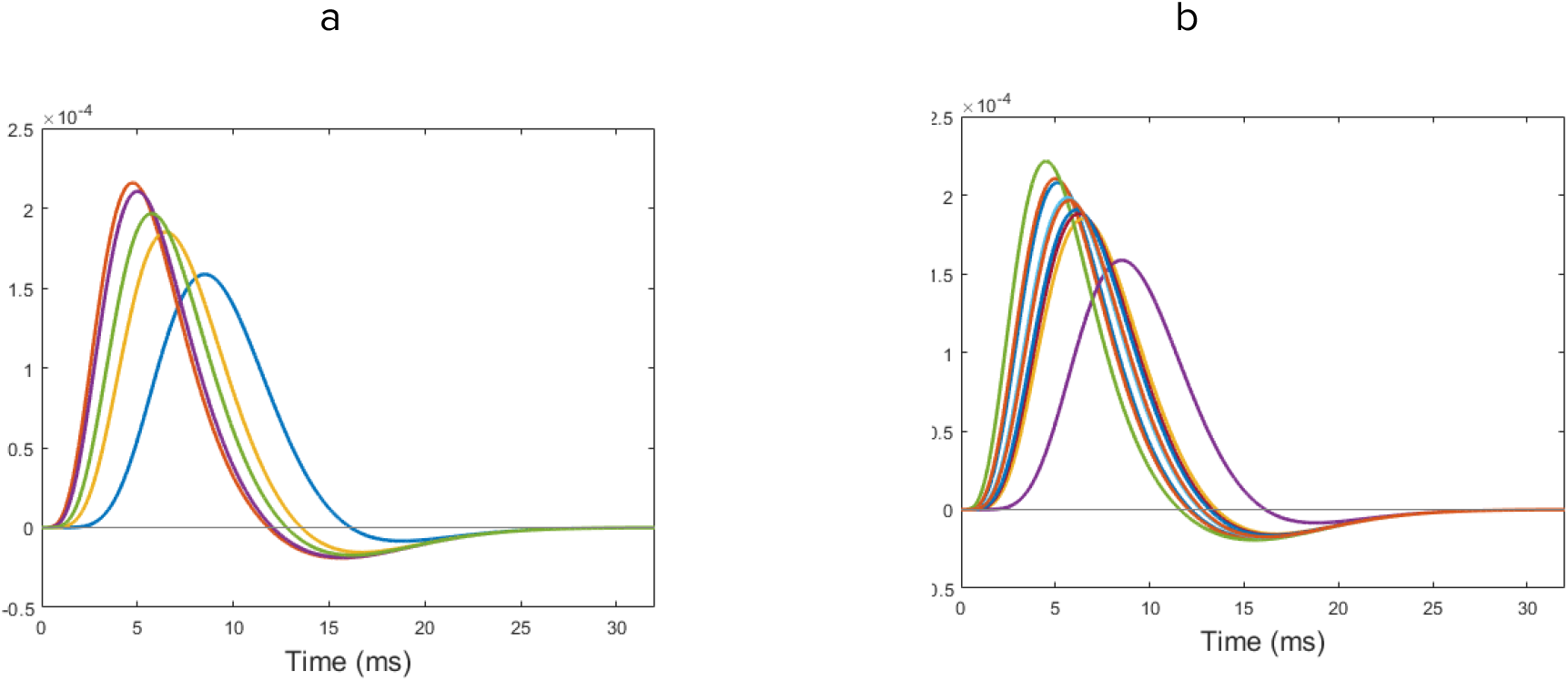
HRFs generated for model A (5-variable model - panel a) and for models B and C (9-variable models - panel b).

**Figure 3.**
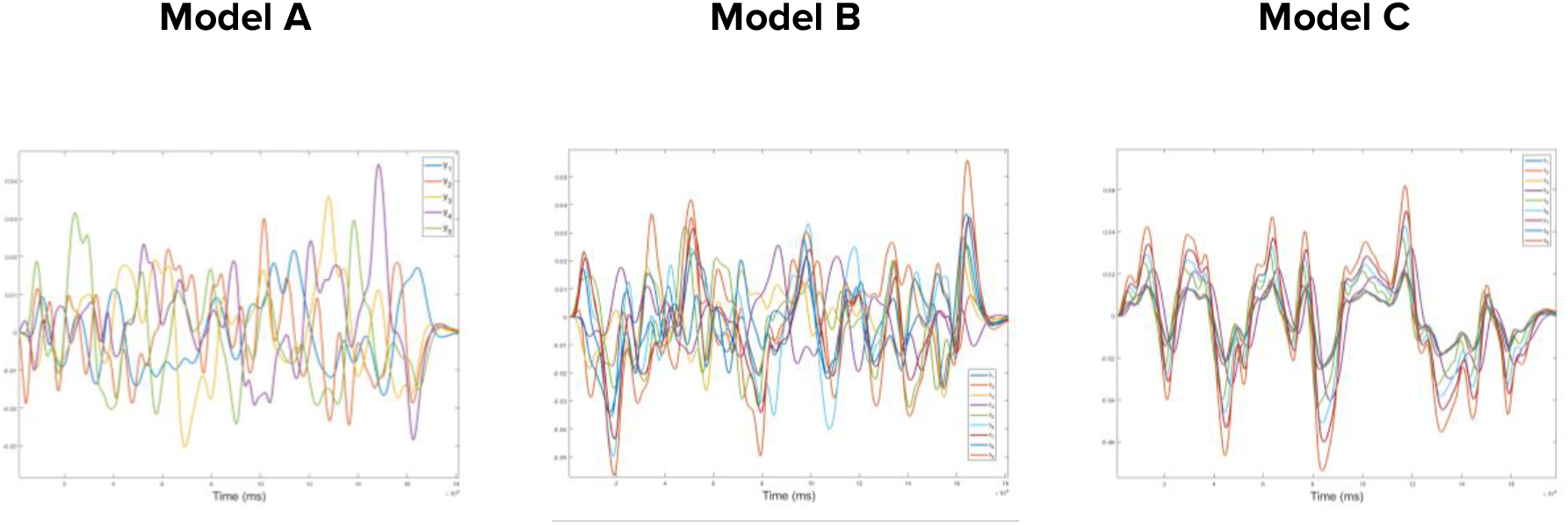
Output of the generative models A, B and C after convolution with the HRFs.

### 2.5. Parameter exploration

#### 2.5.1. Segmentation of generated data

Exploring the differences between connectivity during task and rest is important to characterize reproducibility and replicability of connectivity measures. Previous studies discussed reliability of functional connectivity estimation considering shorter or longer functional MRI scans (for a comprehensive study, see (Shah et al., 2016)).

To this end, we explored a range of values for the number of blocks ([2, 4, 8, 13, 15, 17, 25, 30, 40]) and the number of points per block ([10, 20, 30, 40, 60, 80, 100, 160, 200]). Additionally, to minimize non-stationarity, data were demeaned and detrended.

#### 2.5.2. Signal to noise ratio

In order to study the impact of SNR (conceptualized by comparing the signal of interest with the background noise) on the efficiency of connectivity estimation, we considered three different SNR values: 1000, 200 and 1. In the context of fMRI, SNR has been reported ranging from 4.42 to 280 (Welvaert and Rosseel, 2013).

#### 2.5.3. Subsampling

Previous studies have identified limitations in the estimation of GC associated with sub-sampled time series. In this sense, as evidence showed that subsampling neural processes can distort causality (Barnett and Seth, 2017), we investigated the relation between sampling frequency (Fs) and causality using synthetic data at three sampling rates: 1000 Hz - that matches the generative model sampling, 250 Hz - subsampling with a factor of 4 which approximates the sampling rate of EEG data, and 0.5 Hz - subsampling with a factor of 2000 corresponding to fMRI data with TR of 2 seconds.

### 2.6. Practical implementation

We defined a two step approach (Figure 4) implemented in MATLAB 2016b (Mathworks Inc., USA). In the first step, we explored the combination of estimation method and statistical validation approach, comparing MVGC (MATLAB implementation described in (Barnett and Seth, 2014)) and SS-GC (Barnett and Seth, 2015), coupled with three different approaches for statistical validation: a) time reversed testing, b) PS, and c) theoretical statistical test for MVGC (as in (Barnett and Seth, 2014)). For both test a) and b) a two-sided *t*-test was used to assess whether net connectivity scores were significantly different from zero across repetitions. More specifically, in test a) time-reversed data was built by time inversion testing of each block of the respective MVAR model created, while in test b) the *t*-test was performed on the differences of scores obtained on original and temporally permuted data. In both cases the resulting *t*-scores were converted into *z*-scores. Throughout the paper, connections were reported as significant if the *p*-value associated with the *z*-score falls below a statistical significance *α*=0.05 corrected with family-wise error (FWE) method, to address multiple comparison limitations.

**Figure 4.**
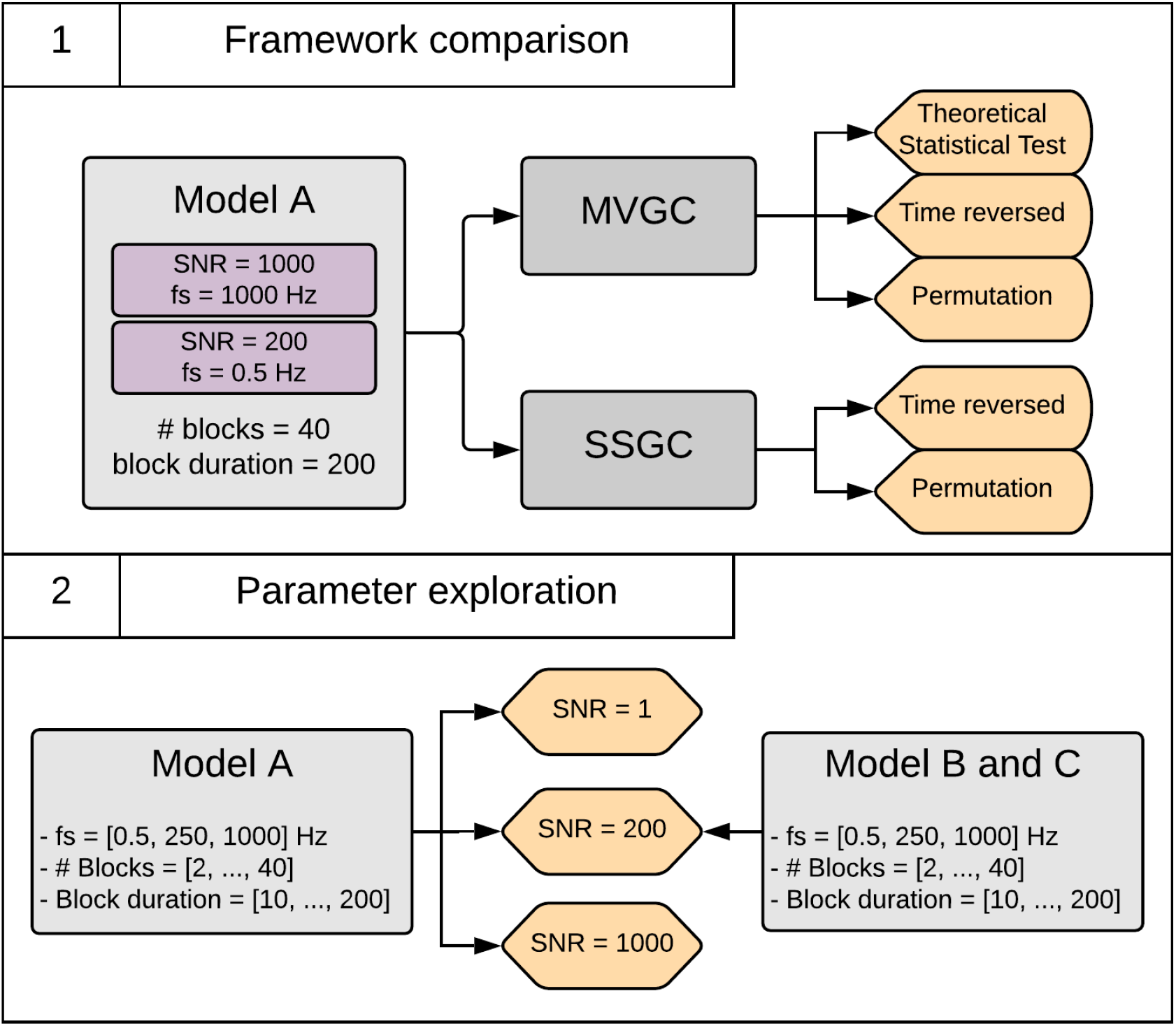
Two step approach considered: 1 - Framework comparison, testing a set of parameters for different metrics of GC and their statistical implementation were tested. 2 - Framework Exploration: number of blocks, duration of blocks, sampling frequency, SNR and number of variables for Model A, B and C.

We assumed a model order of 1, due to the short lag considered in the generative model, and as a methodological standard across the different tests (Schmidt et al., 2016; Zhang et al., 2016).

We used two datasets: i. fine time scale (matching the generative model sampling) and high SNR, and ii. subsampled time series (0.5 Hz) with lower SNR of 200 (within the range described in (Welvaert and Rosseel, 2013)).

Once established the optimal framework (method and validation), we characterized its applicability with a wide range of parameters (number of blocks and points, sampling rate, SNR, number of variables and their mean correlation).

#### 2.6.1. Model validation and Outcome metric

Across the study, validation tools were used to assess robustness of the results.

To assess the fit of the created models compared to the synthetic datasets we used consistency analysis (Rodrigues and Andrade, 2014). This quality measure of the model is important to determine possible confounds related with poor modelling. In accordance with (Ding et al., 2000; Seth, 2010), we used a consistency threshold of 80% to define valid VAR models. Consistency explores the fit of the model analyzing the correlation structure of the original data segments with model generated data segments:

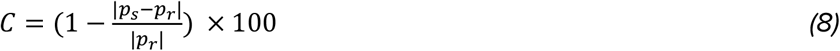

where *p*_*r*_is the correlation vector of the generated data and *p*_*s*_is the correlation vector of the simulated data obtained via the VAR model.

The F1-score, our main outcome measure for each test, is the harmonic mean of the precision and recall and was obtained by comparing the true label of the causal influences in the generative model with the outcome statistically validated connections of the tested framework.

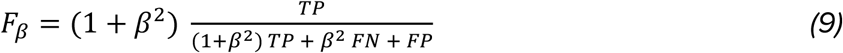

where, *β* is 1, TP represents the true positives, FN false negatives (type II errors) and FP the false positives (type I error). As an example, TP would represent a connections between two variables of the MVAR model that was statistically validated by one of the frameworks studied and would match with the ground truth label of the MVAR model.

## 3. Results

First, we explored different combinations of GC methods and statistical frameworks. Once the best combination/framework was established, we analyzed different parameters such as the number of blocks, number of points per blocks, sampling rate, SNR, number of variables of the network and their mean correlation.

### 3.1. Finding the optimal GC framework / Comparison of GC frameworks

In order to provide an overview of the performance of each framework, the F1-scores are presented in Table 1. Results concern the application to synthetic data of a 5-node network, with high SNR (1000) and without subsampling (equivalent to a 1000 Hz signal).

**Table 1.** F1-scores (a = 0.05 with FWE correction) for the SNR 1000, 5 var, 1000Hz model. SS-GC with time reversed testing, SS-GC with PS, MVGC with the theoretical Statistical Test, MVGC with time reversed testing and MVGC with PS. The acronym n.d. means that F1-score was not defined.

| GC method | Statistical test | F1-score |
| --- | --- | --- |
| SS-GC | Time reversed | 0.67 |
|  | PS | 0.16 |
| MVGC | Theoretical | 0.43 |
|  | Time reversed | n.d. |
|  | PS | 0.34 |

The best results were obtained with the SS-GC method and statistical validation based on time reversed testing, with an F1-score of 0.67.

Table 2 represents the F1-scores regarding the application to synthetic data of a 5-node network, with lower SNR of 200 and coarser subsampling, factor of 2000 (equivalent to a 0.5 Hz signal).

**Table 2.** F1-score values (a = 0.05 with FWE correction) for the SNR 200, 5 MVAR, 0.5 Hz model. SS-GC with time reversed testing, SS-GC with PS and MVGC with the theoretical Statistical Test. The acronym n.d. means that F1-score was not defined.

| GC method | Statistical test | F1-score |
| --- | --- | --- |
| SS-GC | Time reversed | 0.53 |
|  | PS | n.d. |
| MVGC | Theoretical | 0.24 |
|  | Time reversed | n.d. |
|  | PS | n.d. |

The framework presenting higher F1-score for this dataset was also the combination of SS-GC with time reversed testing (F1-score of 0.53). Generally, the methods present worse results compared to the previous dataset.

Missing values - *n.d.* from Table 1 and Table 2, result from intrinsic instability of the modeled time series, i.e. the VAR model was unable to converge, since the spectral radius of the VAR was higher than one (see for a detailed explanation of spectral radius calculation Barnett, et. al 2014).

### 3.2. Effects of *dataset parameters* on causality estimation

To systematically explore the differences associated with dataset parameters, we performed a parametric evaluation of the framework SS-GC with time reversed testing. An overview of the best configuration is presented in Table 3.

**Table 3.**
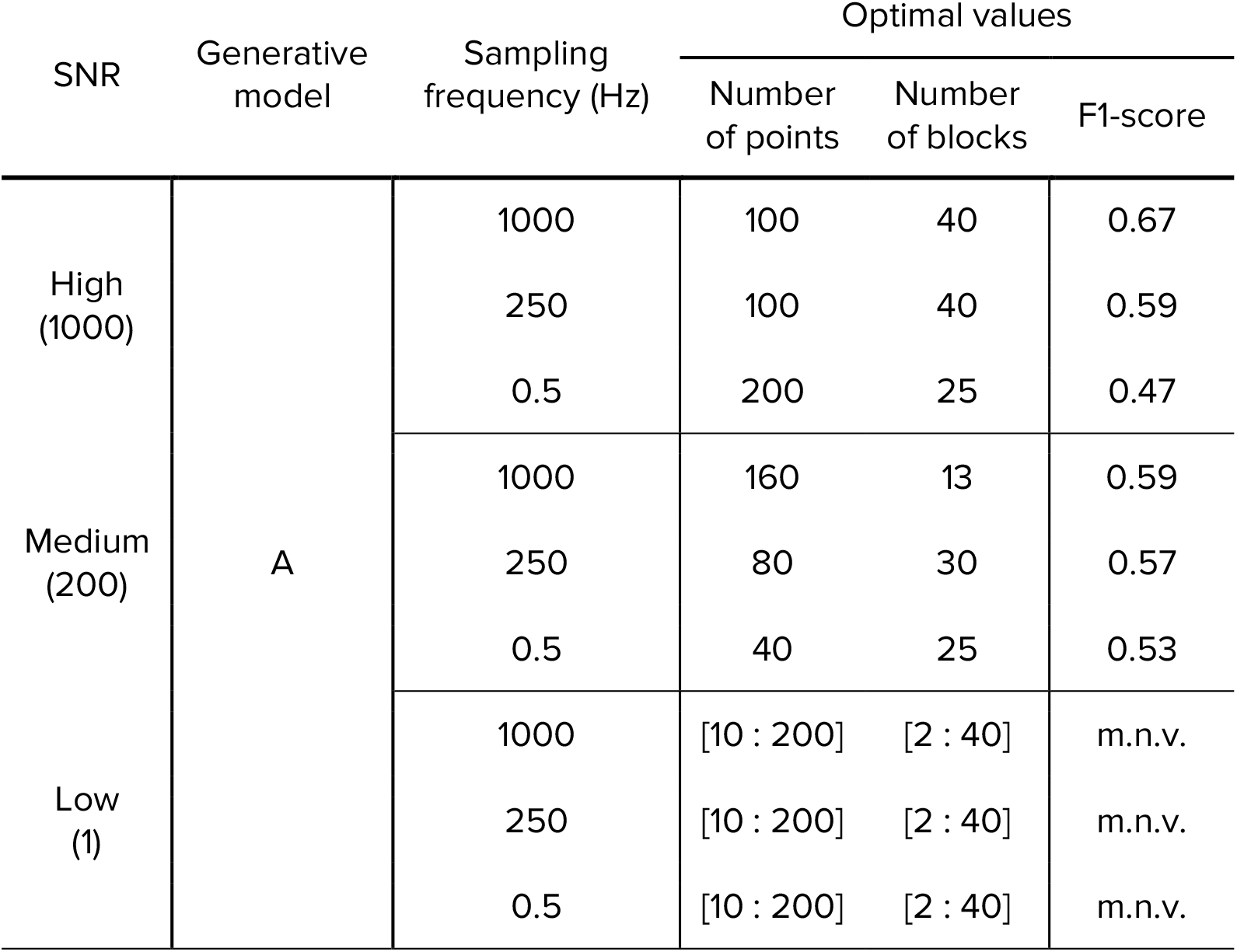
Framework exploration of generative data model A. Maximum F1-score (a=0.05 with FWE correction) with the corresponding values for number of points,number of blocks, variation of SNR and sampling. m.n.v. - model not valid, consistency of the model lower than 80%.

#### 3.2.1. Number of blocks and points per block

To look at the effect of the size and number of blocks in block-partitioned datasets (reflecting task-based paradigms), we considered 2 to 40 blocks and 10 to 200 samples per block. The generative model considered to create the synthetic datasets was based on the 5-node network (model A), with SNR of 1000 and no subsampling (Figure 5.a).

**Figure 5.**
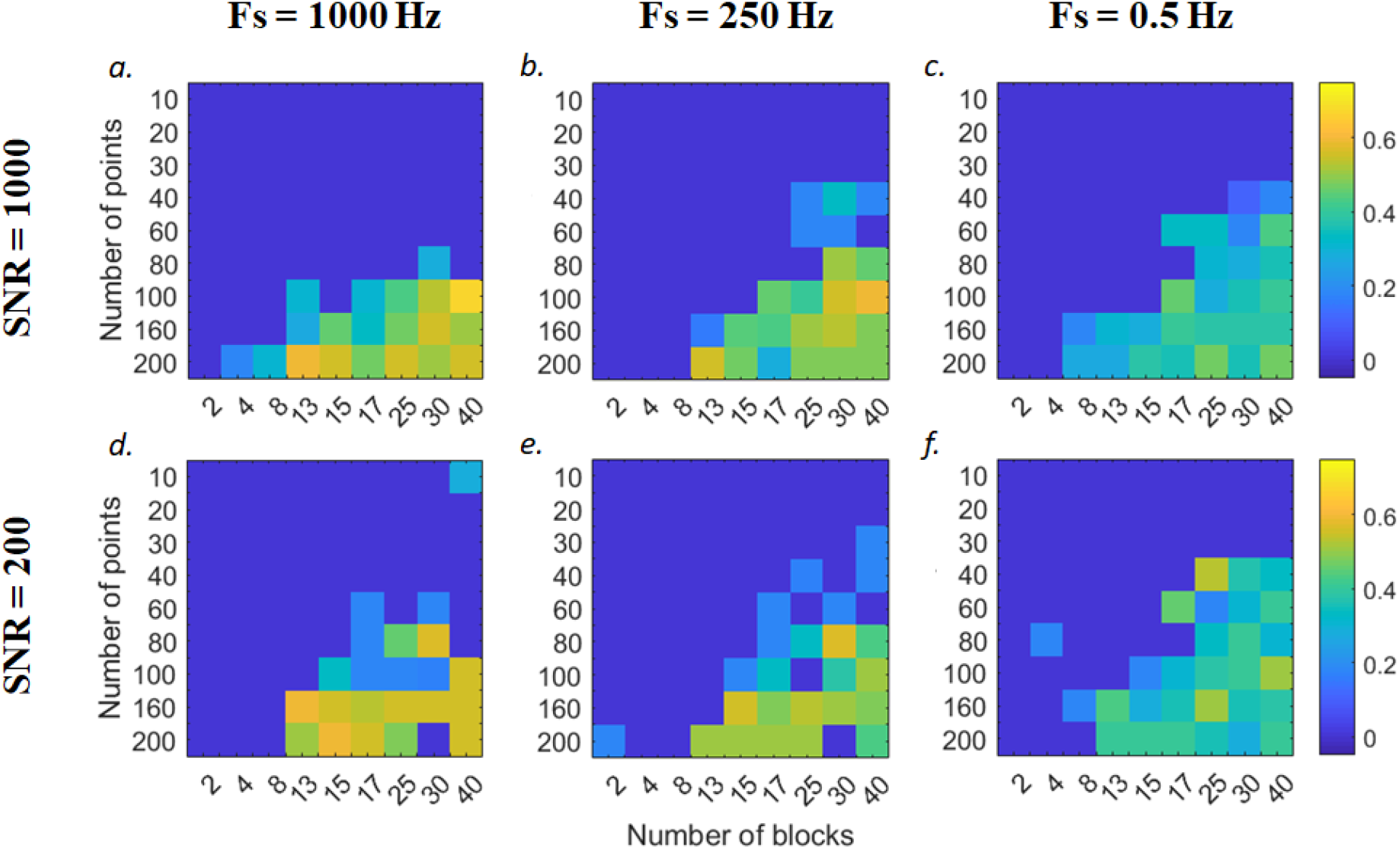
*Distribution of the F1-score values (a=0.05 with FWE correction) across the number of points and number of blocks. Values are depicted for the implementation of SS-GC with* time reversed testing *to the model A. Each shows data with different sampling, 1000 Hz; 250 Hz & 0.5 Hz. The top line represents the model A with SNR = 1000 and the bottom line represents the model A with SNR = 200.*

The results suggest that an increase in the number of samples available to compute GC, i.e. increase of number of blocks and samples per block, improves the ability to correctly identify connections (at least 800 available samples - 4 blocks, 200 samples per block). The best result is obtained with 4000 samples (40 blocks, 100 samples per block), with an F1-score of 0.67.

#### 3.2.2. Sampling frequency

Figure 5.b presents F1-score results for the two subsampled datasets. The best F1-score for the 250 Hz dataset was 0.59, obtained with 40 blocks and 100 points per block, while for the 0.5 Hz dataset the maximum F1-score was 0.47, obtained with 25 blocks and 200 points per block.

#### 3.2.3. Variability based on SNR / consistency analysis

To examine the effects of SNR on GC estimation, we analyzed F1-scores as function of different SNRs (1, 200, and 1000). The synthetic data considered was created based on generative model A. Figure 5.c presents the F1-scores of the different SNR and subsampling factor of 2000 (equivalent to 0.5 Hz datasets).

As expected, the best result was obtained with the highest SNR and without subsampling. Subsampling and lower SNRs impact the performance of GC estimates. Consistency analysis results for lower SNR (as shown in Figure 6), presented values below 80%, meaning that for noisy signals, the reconstructed model is unable to recover the original characteristics of the data.

**Figure 6.**
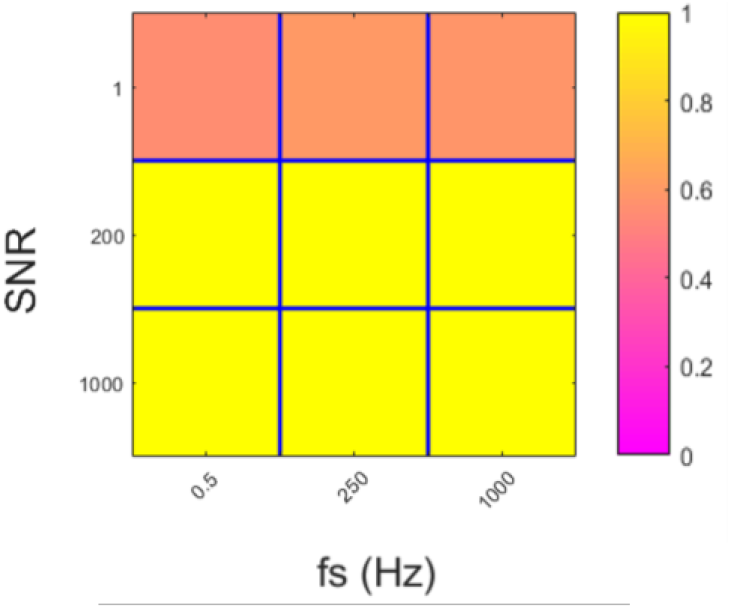
Consistency values for different model parameters. Values of consistency across blocks with 160 points; for model A for each different sampling frequency and each SNR value. The colorbar gives the mean value of consistency.

#### 3.2.4. Number of variables in the generative model and its mean correlation

Two different generative models, B and C, were used to explore the effect of the mean correlation between nodes in connectivity estimation. The F1-score results (Figure 7) were obtained based on the models with SNR = 200.

**Figure 7.**
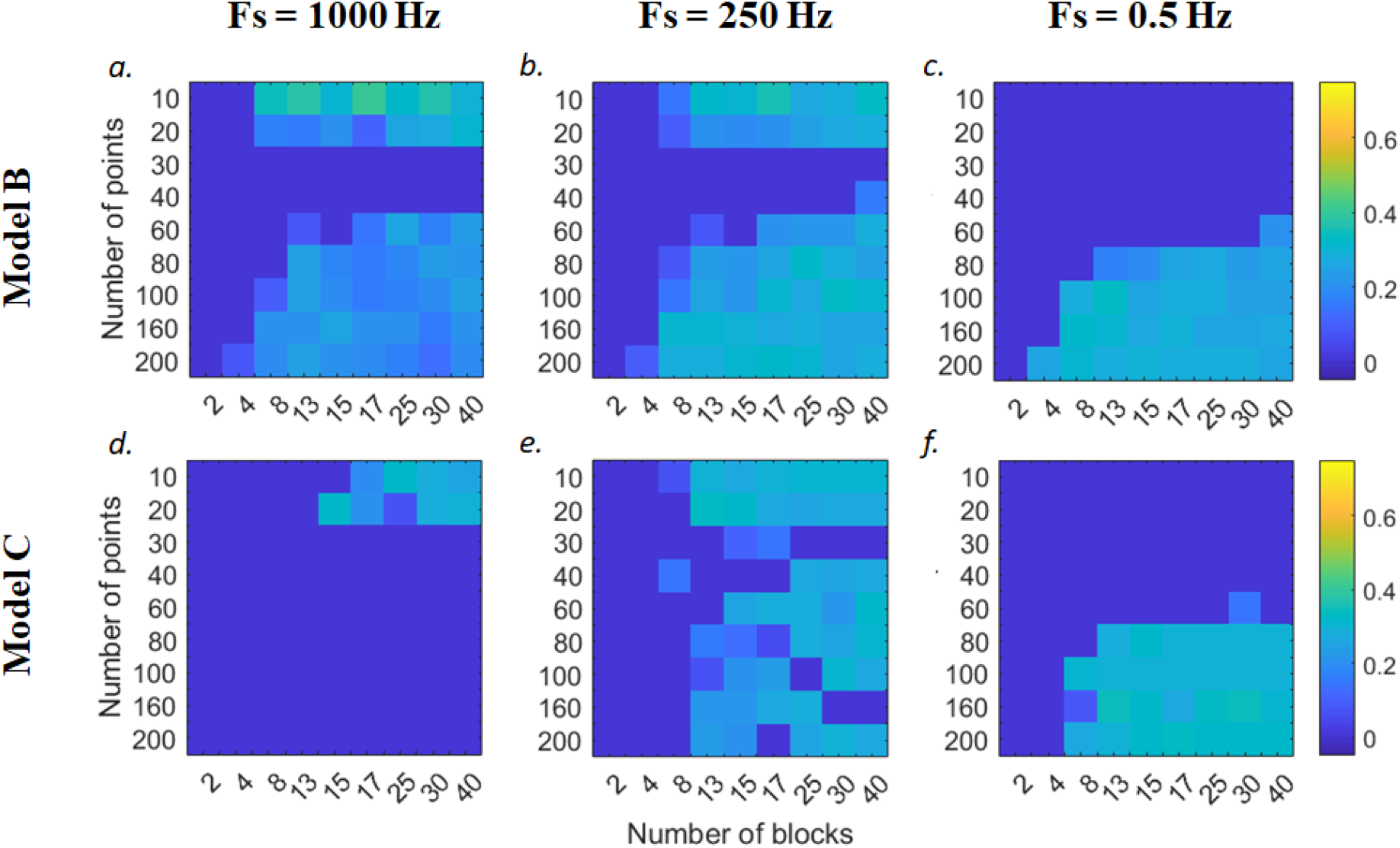
*Distribution of the F1-score values (a=0.05 with FWE correction) across the number of points and number of blocks. There are depicted values for the implementation of SS-GC with* time reversed testing *(SNR = 200). Each column shows data with different sampling, 1000 Hz; 250 Hz & 0.5 Hz. The top line represents the model B. The bottom line represents the values for the model C.*

F1-scores obtained for model B seem to suggest that stronger subsampling impacts the identification of connectivity patterns. On the opposite, there is no impact of subsampling on the best result considering the generative model C. An overview can be depicted on Table 4.

**Table 4.** Framework exploration of generative data model B and C. Maximum F1-score (a=0.05 with FWE correction) with the corresponding values for number of points and number of blocks and sampling. SNR was constant and equal to 200. A consistency higher than 80% was verified.

| SNR | Generative model | Sampling frequency (Hz) | Optimal values |  |  |
| --- | --- | --- | --- | --- | --- |
|  |  |  | Number of points | Number of blocks | F1-score |
| Medium (200) | B | 1000 | 10 | 17 | 0.41 |
|  |  | 250 | 10 | 17 | 0.36 |
|  |  | 0.5 | 100 | 13 | 0.33 |
|  | C | 1000 | 20 | 13 | 0.32 |
|  |  | 250 | 20 | 4 | 0.30 |
|  |  | 0.5 | 200 | 13 | 0.33 |

Considering the 0.5 Hz subsampling, the results show that there is a slight decrease in the F1-score (F1-score maximum for model B = 0.33 < F1-score max for model A = 0.53, respectively - see Table 4) and require a similar number of blocks and number of points per block.

## 4. Discussion

GC has been explored as a measure of directed functional connectivity in neuroimaging data aiming to investigate effects, rather than underlying mechanisms (Barrett and Barnett, 2013). Nonetheless, its application to different neuroimaging modalities has raised a significant number of technical and experimental challenges.

The validity of analysing GC, particularly in fMRI data, is still a matter of debate. Previous studies have discussed the influence of the hemodynamic response latency (Seth et al., 2013), SNR (Luo et al., 2011; Seth et al., 2013) and subsampling (Rodrigues and Andrade, 2014; Wen et al., 2013). In general, these studies suggest that such parameters contribute as confounds, increasing the risk of detecting spurious connectivity (David et al., 2008; Smith et al., 2012).

In an attempt to address these challenges, novel approaches have been recently proposed both in the context of model estimation and statistical validation (Barnett and Seth, 2017, 2015; Faes et al., 2017; Haufe et al., 2013; Solo, 2015; Winkler et al., 2016).

In this paper, we address SS-GC combined with time reversed testing, compare it with MVGC approach, and test its performance with a number of technical and experimental parameters. Ultimately, we aimed to explore the possible shortcomings of applying GC to fMRI data, by synthetically creating datasets with a set of parameters which could characterize its applicability. In this sense, our objective is also to characterize the optimal set of parameters (in terms of number of blocks and points per block) for each scenario.

To this end, we considered three generative models (varying in number of nodes and their connection, models A, B and C) convolved with confounding HRFs (Seth et al., 2013). Based on these models, we simulated a number of datasets with different number of samples (time series length), data partition in blocks, downsampling (sampling frequency of the signal), and SNR. This allowed us to compare GC in different data architectures (while considering the structure of neuroimaging datasets) and explore the applicability of the causality metrics in fMRI scenarios over different timescales. The conceptualization of BOLD as a representation of the millisecond processes involved in neuronal activity has been previously addressed by (Logothetis et al., 2001), that showed that BOLD represents a low pass filter version of LFPs and that significant physiological patterns can be found in different time scales (e.g. second).

We measured the ability of the computational frameworks to correctly identify synthetic connections, by obtaining F1-scores of the comparison between the true and predicted connections.

### 4.1. Optimal combination of GC estimation method and statistical framework

The framework combining SS-GC and time reversed testing presented the best results considering the dataset obtained with a 5-node network, with high SNR (1000) and without subsampling (equivalent to a 1000 Hz signal).

SS modelling explicitly includes observational noise in the model (Barnett and Seth, 2015; Solo, 2016; Sommerlade et al., 2015). The parameters are estimated with reduced bias, contributing to a simpler, more computationally efficient and numerically stable solution. Conversely, MVGC performance was greatly challenged by the addition of noise to the data.

Our results also suggest that time reversed testing outperforms PS in both GC methods. The limitations of PS were previously addressed by (Haufe et al., 2013), demonstrating that, using simulated EEG data, permutation testing led to the identification of spurious connectivity.

(Winkler et al., 2016) explored the empirical and theoretical grounds of time reversed testing. They outlined its behavior on additive noise models, and the impact of downsampling and temporal aggregation in the presence of causal interactions. The authors concluded that time reversed testing may robustly reject causal interpretations for mixtures of non-interacting sources, which may arise in realistic settings due to volume conduction and subsampling (EEG and fMRI scenarios, respectively).

### 4.2. Effect of dataset parameters on causality estimation

#### 4.2.1. Number of points and blocks

The number of data points and its organization in blocks showed to have a clear impact on the performance of the framework. Here, we explored their empirical boundaries; the best results (considering the generative model A) were found between 13 and 40 blocks, and 40 and 200 points.

(DSouza et al., 2017) showed that both a shorter TR and a longer scan duration improve GC connectivity estimation. The authors advise that if one of the two is limiting in the study protocol, the other should be varied (maximizing the number of samples) to improve brain connectivity analysis results (Cheung et al., 2010). In addition, (Bastos and Schoffelen, 2016) show that there may be a tendency to overestimate the connectivity in the dataset with the smallest sample size.

#### 4.1.2. Subsampling

The discretization of continuous data processes and the effect of downsampling in the estimation of GC was also studied. Our results suggest that increasing the subsampling factor, the detectability decreases. Considering the model A, we observe a decrease in the best F1-score associated with the increase in the subsampling factor, even considering different SNR. This effect is not noticeable for model C (more complex and with stronger correlations between nodes) - the F1-scores are lower overall but comparable considering different subsampling factors.

(Winkler et al., 2016) concluded that there is a decrease in the true positive rate associated with the increasing factor of downsampling. Our results suggest a similar pattern.

Previous studies explored the complex relation between subsampling and GC estimation. (Zhou et al., 2014) demonstrated that time series obtained by using different sampling interval lengths lead to great differences in GC values. Importantly, they found that a finer sampling did not result in more reliable GC values for causal interpretation. (Barnett and Seth, 2017) concluded that detectability tends to decrease as the sample time interval increases beyond causal delay times, and suggests the existence of “black spots”, where detectability becomes difficult or unfeasible, and then recovers up to a “sweet spot”. In this sense, the authors show that downsampling may even improve detectability.

#### 4.1.3. SNR

One of the main challenges regarding the estimation of GC is that measurement noise can degrade the causal relation between time series and its detectability as well as lead to the detection of spurious connections. The signals measured by electrophysiology and fMRI contain a poorly known mixture of signal-of-interest and “noise” (Winkler et al., 2016).

Our results showed that the SNR has a strong impact on the framework performance. The results confirm that for the same model and sampling frequency, greater the SNR, greater the F1-scores overall. Strikingly, in the case of SNR=1, the reconstructed models were unable to recover the original characteristics of the data.

(Bastos and Schoffelen, 2016; Nalatore et al., 2014, 2007) studied how GC can be corrupted by differences in SNR. Both addressed bivariate models with equal theoretical connectivity strengths, added noise to one of the variables, and studied its effect over the connectivity estimates. The conclusion was that the additional noise on one of the variables had weakened the predictive power of the other variable, causing an apparent asymmetry in the directionality. Asymmetries in GC that are driven by SNR differences, which create spurious connectivity, have been defined as “weak asymmetries” (Haufe et al., 2012) and studied with time reversed testing (Winkler et al., 2016). The authors concluded that time reversed testing reduces this effect. (Barnett and Seth, 2015; Sommerlade et al., 2015) also addressed the SNR challenge considering the SS model, trying to separate the noise component. The conclusions were similar: the stronger the impact of the noise, the more difficult it was to recover the connectivity structure of the generative model and to prevent inference of wrong connectivity.

#### 4.1.4. Number of variables and their mean correlation

Increasing the complexity of the model (e.g. model order or number of network variables) may pose additional computational and analytical challenges. For example, the condition N - p ≥ D, where N represents the number of points, p the model order and D the number of network variables, has to be fulfilled (Seth, 2010).

To assess its impact in the proposed technique, we compared i. the effect of the number of variables, and ii. the effect of the magnitude of the mean pairwise correlation.

i. The increase in the number of nodes of the network reduced F1-score values overall, irrespective of the subsampling factor. (Rodrigues and Andrade, 2014) verified that the sensitivity of the causality measures degrades very rapidly with the increase in the number of nodes, if not followed by the increase in the time series length, an observation that our tests reinforce. A new approach, large-scale GC (lsGC), estimates multivariate dFC in datasets with high number of nodes, incorporating an embedded dimension reduction (DSouza et al., 2017; Pester, 2013; Pester et al., 2015; Schmidt et al., 2016, 2014). Multiple tests showed that complex network structures could be captured by lsGC in both simulated and resting-state fMRI data, even when the number of variables greatly exceeded the number of temporal samples. lsGC addresses this challenge and is especially promising for neuroimaging studies (e.g. fMRI), where the number of voxels vastly exceeds the number of samples.

ii. We found that higher mean pairwise correlation between network variables also limits connectivity estimation performance. This was also identified as a major issue for GC measures in other studies (Barnett and Seth, 2014; Haufe et al., 2013), namely for EEG where naturally occurring highly correlated data complicates application of GC (Flamm et al., 2013).

### 4.2. Limitations

The connections in the synthetic data were simulated considering a discrete process with intervals of 20 ms, which is in line with the literature on neural delays (Schippers et al., 2011). As so, the time-scale of the generative model, which only mimics data from millisecond ‘neural’ activity events to the second ‘neuroimaging’ temporal scales (i.e. once downsampled), should be considered a methodological restriction. In this sense, the predictions of the causal relations could only aim to uncover the network topology depending on those specific neural delays, even though we do not exclude the possibility of real data having longer or shorter neural delays.

The estimation of the GC model requires the definition of the model order (p). An underestimated (or overestimated) model order may influence the results if the process characteristics are not being properly reflected by the model (Bressler and Seth, 2011; Schmidt et al., 2016). Nonetheless, we chose to define a model order of 1 since it relates to the timescale of measured physiological data. A higher p would add much delayed information to the model and lead to inflated Type I and Type II error rates (Barnett and Seth, 2011), while increasing exponentially the computational effort and its feasibility. We have also empirically tested the models evaluating the statistical consistency (above the limit of 80% as demonstrated in (Seth, 2010)).

### 4.3. Future Work

Future work should consider increasing the empirical foundations of SS-GC, exploring the boundaries here described to discriminate optimal scenarios; our results showed promise of the application to synthetic data with specific temporal resolution, which precludes naive application of the metric to fMRI data in general. Nevertheless, the validity of the algorithms in real-world scenarios, in which the ground truth is unknown, requires the determination of reliability. Future research should address the assessment of the reliability of these methods for the test-retest convergence (Bielczyk et al., 2018).

### 4.4. Conclusions

The study presents an innovative approach towards the validation of SS-GC in the context of generative models. By coupling it with time reversed testing, we were able to estimate reliable causal relations in a number of synthetic datasets.

While the fMRI scenario is perhaps one of the most challenging for causality measures, our framework showed promising results regarding the detection of incorrect connections, emphasizing the importance of having a high number of samples per condition and a high SNR. Recent technological advances with ultra-fast/multiband MRI sequences (Feinberg and Setsompop, 2013) and higher magnetic field (Duggento et al., 2016) pave the way for improved neuroimaging datasets.

In conclusion, our results offer an empirical demonstration of the possible applicability boundaries of SS-GC in neuroimaging scenarios.

## Acknowledgements

We acknowledge the Portuguese Science Foundation (FCT) for financial support through “Projecto Operacional Regional do Centro”-BIGDATIMAGE: CENTRO-01-0145-FEDER-000016 and MEDPersyst POCI-01-0145-FEDER-016428; Contract grant sponsor: FCT; Contract grant number: UID/NEU/ 04539/2013; UID/4950/2020, Contract grant sponsor: COMPETE; Contract grant number: POCI-01-0145-FEDER-007440, PTDC/PSI-GER/30852/2017 and the European Commission, H2020-SC1-2016-2017, STIPED.

## Supplementary data

### Mathematical models

We describe the model mathematically supporting dataset A in Equation S.1:

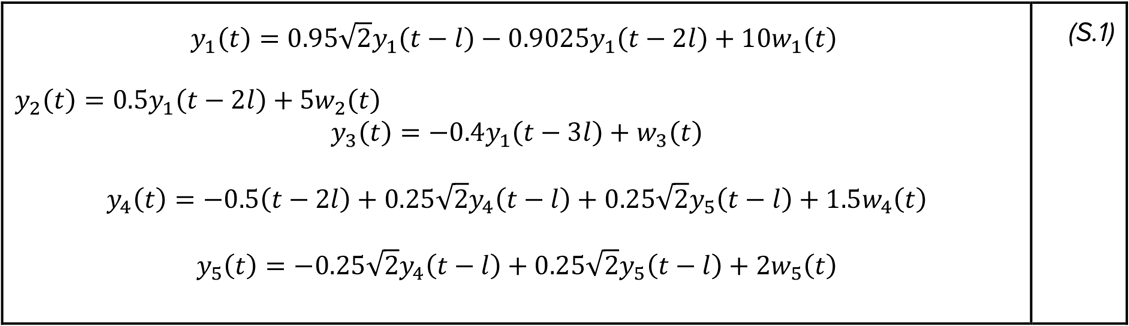

where *y*_*n*_(*t*), *n* = 1, … , *N*, represents the value of the variable *y* at instant *t*, *l* is the lag between two consecutive samples of the AR model and *w* _*n*_(*t*) represents zero-mean uncorrelated white processes with identical variance.

Dataset B is mathematically described by Equation S.2:

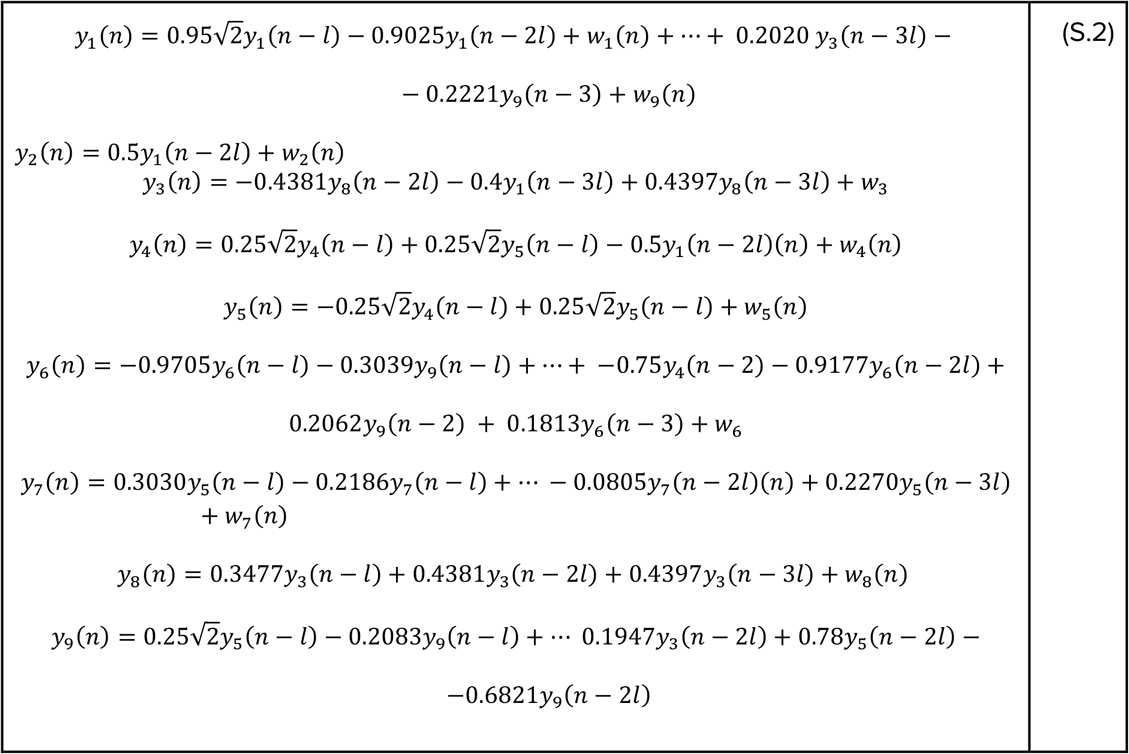

The last model, dataset C is also composed by 9 variables but presents higher correlation coefficients between variables and is mathematically defined by equation S.3:

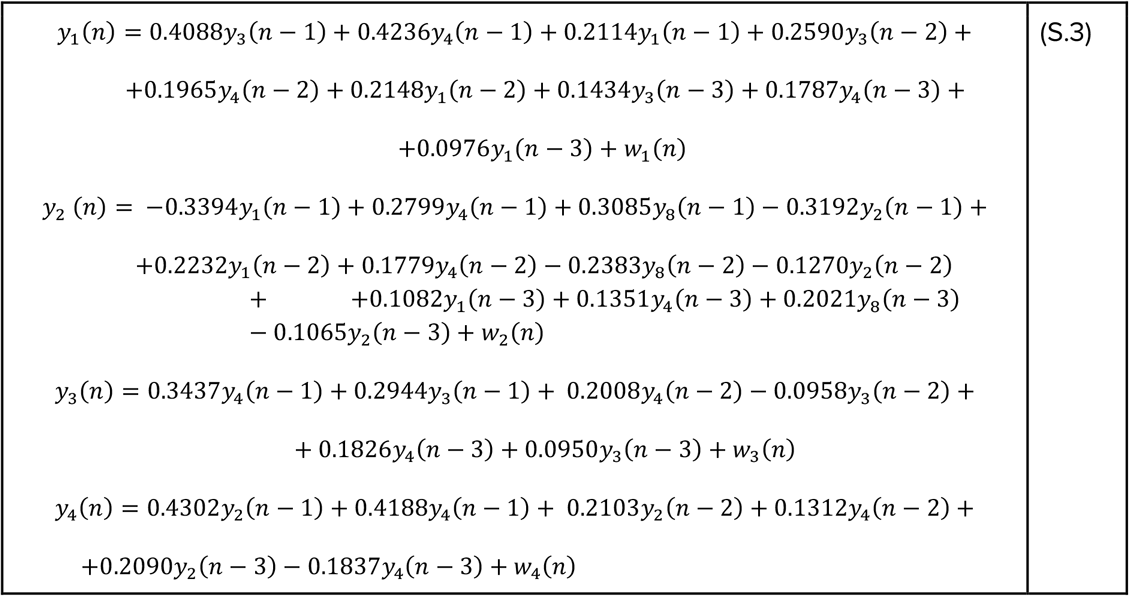

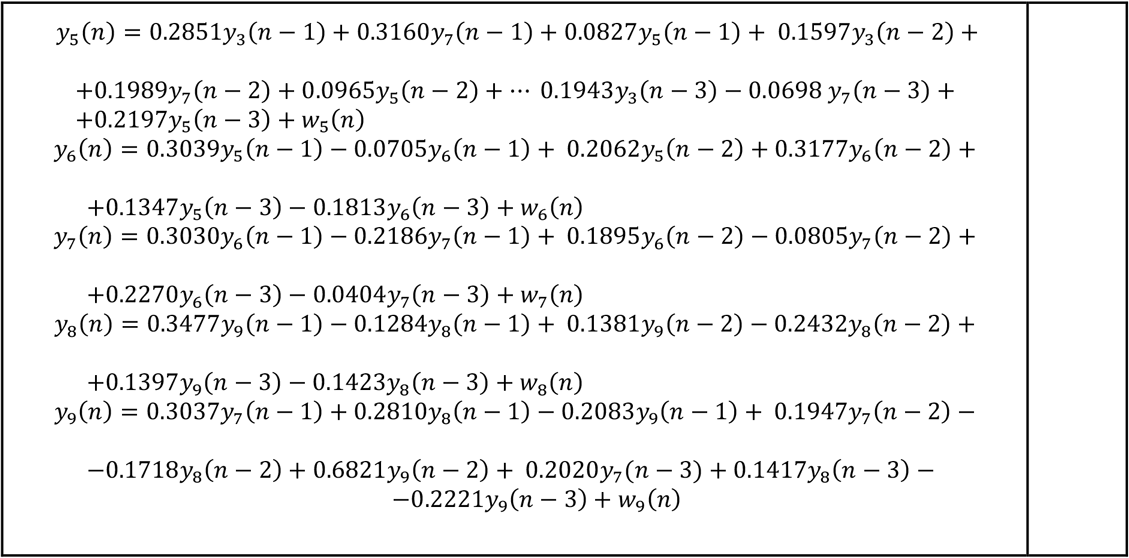

## References

Anderson, M.J., Robinson, J., 2001. Permutation tests for linear models. Aust. N. Z. J. Stat. 43, 75–88. https://doi.org/10.1111/1467-842X.00156

Baccalá, L.A., Sameshima, K., 2001. Partial Directed Coherence: a new Concept in Neural Structure Determination. Biol. Cybern. 84, 463–474.

Barnett, L., Barrett, A.B., Seth, A.K., 2018. Solved problems for Granger causality in neuroscience: A response to Stokes and Purdon Lionel. Neuroimage. https://doi.org/10.1016/j.neuroimage.2018.05.067

Barnett, L., Seth, A.K., 2017. Detectability of Granger causality for subsampled continuous-time neurophysiological processes. J. Neurosci. Methods 275, 93–121. https://doi.org/10.1016/j.jneumeth.2016.10.016

Barnett, L., Seth, A.K., 2015. Granger causality for state-space models. Phys. Rev. E 91, 1–5. https://doi.org/10.1103/PhysRevE.91.040101

Barnett, L., Seth, A.K., 2014. The MVGC multivariate Granger causality toolbox: a new approach to Granger-causal inference. J. Neurosci. Methods 223, 50–68. https://doi.org/10.1016/j.jneumeth.2013.10.018

Barnett, L., Seth, A.K., 2011. Behaviour of Granger causality under filtering: theoretical invariance and practical application. J. Neurosci. Methods 201, 404–19. https://doi.org/10.1016/j.jneumeth.2011.08.010

Barrett, A.B., Barnett, L., 2013. Granger causality is designed to measure effect, not mechanism. Front. Neuroinform. 7, 6–7. https://doi.org/10.3389/fninf.2013.00006

Barrett, A.B., Murphy, M., Bruno, M.-A., Noirhomme, Q., Boly, M., Laureys, S., Seth, A.K., 2012. Granger causality analysis of steady-state electroencephalographic signals during propofol-induced anaesthesia. PLoS One 7, e29072. https://doi.org/10.1371/journal.pone.0029072

Bastos, A.M., Schoffelen, J.-M., 2016. A Tutorial Review of Functional Connectivity Analysis Methods and Their Interpretational Pitfalls. Front. Syst. Neurosci. 9, 1–23. https://doi.org/10.3389/fnsys.2015.00175

Bielczyk, N.Z., Uithol, S., van Mourik, T., Anderson, P., Glennon, J.C., Buitelaar, J.K., 2018. Disentangling causal webs in the brain using functional magnetic resonance imaging: A review of current approaches. Netw. Neurosci. 3, 237–273. https://doi.org/10.1162/netn_a_00062

Bressler, S.L., Seth, A.K., 2011. Wiener-Granger causality: a well established methodology. Neuroimage 58, 323–9. https://doi.org/10.1016/j.neuroimage.2010.02.059

Brovelli, A., Ding, M., Ledberg, A., Chen, Y., Nakamura, R., Bressler, S.L., 2004. Beta oscillations in a large-scale sensorimotor cortical network: Directional influences revealed by Granger causality. Proc. Natl. Acad. Sci. 101, 9849–9854. https://doi.org/10.1073/pnas.0308538101

Cheung, B.L.P., Riedner, B.A., Tononi, G., Veen, B.D.Van, 2010. “Estimation of cortical connectivity from {EEG} using state-space models.,” {IEEE} {Transactions} on {Biomedical} engineering 57, 2122–2134.

David, O., Guillemain, I., Saillet, S., Reyt, S., Deransart, C., Segebarth, C., Depaulis, A., 2008. Identifying neural drivers with functional MRI: an electrophysiological validation. PLoS Biol. 6, 2683–2697. https://doi.org/10.1371/journal.pbio.0060315

Ding, M., Bressler, S.L., Yang, W., Liang, H., 2000. Short-window spectral analysis of cortical event-related potentials by adaptive multivariate autoregressive modeling: data preprocessing, model validation, and variability assessment. Biol. Cybern. 83, 35–45. https://doi.org/10.1007/s004229900137

DSouza, A.M., Abidin, A.Z., Leistritz, L., Wismüller, A., 2017. Exploring connectivity with large-scale Granger causality on resting-state functional MRI. J. Neurosci. Methods 287, 68–79. https://doi.org/10.1016/j.jneumeth.2017.06.007

Duggento, A., Bianciardi, M., Passamonti, L., Wald, L.L., Guerrisi, M., Barbieri, R., Toschi, N., 2016. Globally conditioned Granger causality in brain–brain and brain–heart interactions: a combined heart rate variability/ultra-high-field (7 T) functional magnetic resonance imaging study. Philos. Trans. R. Soc. A Math. Phys. Eng. Sci. 374, 20150185. https://doi.org/10.1098/rsta.2015.0185

Faes, L., Nollo, G., Stramaglia, S., Marinazzo, D., 2017. Multiscale Granger causality. Phys. Rev. E 96. https://doi.org/10.1103/PhysRevE.96.042150

Feinberg, D.A., Setsompop, K., 2013. Ultra-fast MRI of the human brain with simultaneous multi-slice imaging. J. Magn. Reson. 229, 90–100. https://doi.org/10.1016/j.jmr.2013.02.002

Flamm, C., Graef, A., Pirker, S., Baumgartner, C., Deistler, M., 2013. Influence analysis for high-dimensional time series with an application to epileptic seizure onset zone detection. J. Neurosci. Methods 214, 80–90. https://doi.org/10.1016/j.jneumeth.2012.12.025

Fox, M.D., 2010. Clinical applications of resting state functional connectivity. Front. Syst. Neurosci. 4. https://doi.org/10.3389/fnsys.2010.00019

Friston, K., Moran, R., Seth, A.K., 2013. Analysing connectivity with Granger causality and dynamic causal modelling. Curr. Opin. Neurobiol. 23, 172–178. https://doi.org/10.1016/j.conb.2012.11.010

Friston K.J., Harrison L. P.W., 2003. Dynamic causal modelling. Neuroimage 19, 1273–1302.

Friston, K.J., Ashburner, J., Kiebel, S.J., Nichols, T.E., Penny, W.., 2007. Statistical Parametric Mapping: The Analysis of Functional Brain Images. Acad. Press.

Gaillard, R., Dehaene, S., Adam, C., 2009. Converging Intracranial Markers of Conscious Access. PLoS Biol. 7. https://doi.org/10.1371/Citation

Ghasemi, M., Mahloojifar, A., 2013. Disorganization of Equilibrium Directional Interactions in the Brain Motor Network of Parkinson’s disease: New Insight of Resting State Analysis Using Granger Causality and Graphical Approach. J. Med. Signals Sens. 3, 69–78.

Granger, 1969. Investigating causal relations by econometric models and crossspectral methods. Econometrica 424–38.

Haufe, S., Nikulin, V. V., Nolte, G., 2012. Alleviating the influence of weak data asymmetries on Granger-causal analyses. Lect. Notes Comput. Sci. (including Subser. Lect. Notes Artif. Intell. Lect. Notes Bioinformatics) 7191 LNCS, 25–33. https://doi.org/10.1007/978-3-642-28551-6_4

Haufe, S., Nikulin, V. V, Müller, K.-R., Nolte, G., 2013. A critical assessment of connectivity measures for EEG data: a simulation study. Neuroimage 64, 120–33. https://doi.org/10.1016/j.neuroimage.2012.09.036

Kadosh, K.C., Luo, Q., de Burca, C., Sokunbi, M.O., Feng, J., Linden, D.E.J., Lau, J.Y.F., 2016. Using real-time fMRI to influence effective connectivity in the developing emotion regulation network. Neuroimage 125, 616–626. https://doi.org/10.1016/j.neuroimage.2015.09.070

Krueger, F., Landgraf, S., Van Der Meer, E., Deshpande, G., Hu, X., 2011. Effective connectivity of the multiplication network: A functional MRI and multivariate granger causality mapping study. Hum. Brain Mapp. 32, 1419–1431. https://doi.org/10.1002/hbm.21119

Logothetis, N.K., J, P., M, A., T, T., A, O., 2001. Neurophysiological investigation of thebasis of the fMRI signal. Nature 1–8.

Luo, Q., Ge, T., Feng, J., 2011. Granger causality with signal-dependent noise. Neuroimage 57, 1422–1429. https://doi.org/10.1016/j.neuroimage.2011.05.054

Mill, R.D., Bagic, A., Bostan, A., Schneider, W., Cole, M.W., 2017. Empirical validation of directed functional connectivity. Neuroimage 146, 275–287. https://doi.org/10.1016/j.neuroimage.2016.11.037

Nalatore, H., Ding, M., Rangarajan, G., 2007. Mitigating the effects of measurement noise on Granger causality. Phys. Rev. E 75, 031123. https://doi.org/10.1103/PhysRevE.75.031123

Nalatore, H., Sasikumar, N., Rangarajan, G., 2014. Effects of measurement noise on Granger causality. Phys. Rev. E - Stat. Nonlinear, Soft Matter Phys. 90, 1–9. https://doi.org/10.1103/PhysRevE.75.031123

Pester, B., 2013. EXPLORING EFFECTIVE CONNECTIVITY BY A GRANGER CAUSALITY APPROACH WITH EMBEDDED DIMENSION REDUCTION. Biomed Tech 58, 24–25. https://doi.org/10.1515/bmt-2013-4

Pester, B., Schmidt, C., Schmid-Hertel, N., Witte, H., Wismueller, A., Leistritz, L., 2015. Identification of whole-brain network modules based on a large scale Granger Causality approach. Proc. Annu. Int. Conf. IEEE Eng. Med. Biol. Soc. EMBS 2015–Novem, 5380–5383. https://doi.org/10.1109/EMBC.2015.7319607

Pievani, M., Filippini, N., Van Den Heuvel, M.P., Cappa, S.F., Frisoni, G.B., 2014. Brain connectivity in neurodegenerative diseases - From phenotype to proteinopathy. Nat. Rev. Neurol. 10, 620–633. https://doi.org/10.1038/nrneurol.2014.178

Rodrigues, J., Andrade, A., 2014. Lag-based effective connectivity applied to fMRI: a simulation study highlighting dependence on experimental parameters and formulation. Neuroimage 89, 358–77. https://doi.org/10.1016/j.neuroimage.2013.10.029

Roebroeck, A., Formisano, E., Goebel, R., 2005. Mapping directed influence over the brain using Granger causality and fMRI. Neuroimage 25, 230–242. https://doi.org/10.1016/j.neuroimage.2004.11.017

Schippers, M.B., Renken, R., Keysers, C., 2011. The effect of intra- and inter-subject variability of hemodynamic responses on group level Granger causality analyses. Neuroimage 57, 22–36. https://doi.org/10.1016/j.neuroimage.2011.02.008

Schmidt, C., Pester, B., Nagarajan, M., Witte, H., Leistritz, L., Wismueller, A., 2014. Impact of multivariate Granger causality analyses with embedded dimension reduction on network modules. Conf. Proc Annu. Int. Conf. IEEE Eng. Med. Biol. Soc. IEEE Eng. Med. Biol. Soc. Annu. Conf. 2014, 2797–2800. https://doi.org/10.1109/EMBC.2014.6944204

Schmidt, C., Pester, B., Schmid-Hertel, N., Witte, H., Wismüller, A., Leistritz, L., 2016. A Multivariate Granger Causality Concept towards Full Brain Functional Connectivity. PLoS One 11, e0153105. https://doi.org/10.1371/journal.pone.0153105

Seth, a. K., Barrett, a. B., Barnett, L., 2015. Granger Causality Analysis in Neuroscience and Neuroimaging. J. Neurosci. 35, 3293–3297. https://doi.org/10.1523/JNEUROSCI.4399-14.2015

Seth, A.K., 2010. A MATLAB toolbox for Granger causal connectivity analysis. J. Neurosci. Methods 186, 262–73. https://doi.org/10.1016/j.jneumeth.2009.11.020

Seth, A.K., Chorley, P., Barnett, L.C., 2013. Granger causality analysis of fMRI BOLD signals is invariant to hemodynamic convolution but not downsampling. Neuroimage 65, 540–555. https://doi.org/10.1016/j.neuroimage.2012.09.049

Shah, L.M., Cramer, J.A., Ferguson, M.A., Birn, R.M., Anderson, J.S., 2016. Reliability and reproducibility of individual differences in functional connectivity acquired during task and resting state. Brain Behav. 6, 1–15. https://doi.org/10.1002/brb3.456

Smith, S.M., Bandettini, P.A., Miller, K.L., Behrens, T.E.J., Friston, K.J., David, O., Liu, T., Woolrich, M.W., Nichols, T.E., 2012. The danger of systematic bias in group-level FMRI-lag-based causality estimation. Neuroimage 59, 1228–1229. https://doi.org/10.1016/j.neuroimage.2011.08.015

Solo, V., 2016. State-Space Analysis of Granger-Geweke Causality Measures with Application to fMRI. Neural Comput. 28, 914–949. https://doi.org/10.1162/NECO

Solo, V., 2015. State Space Methods for Granger-Geweke Causality. arXiv.

Sommerlade, L., Thiel, M., Mader, M., Mader, W., Timmer, J., Platt, B., Schelter, B., 2015. Assessing the strength of directed influences among neural signals: An approach to noisy data. J. Neurosci. Methods 239, 47–64. https://doi.org/10.1016/j.jneumeth.2014.09.007

Stephan, E.K., Roebroeck, A., 2012. NeuroImage A short history of causal modeling of fMRI data. Neuroimage 62, 856–863. https://doi.org/10.1016/j.neuroimage.2012.01.034

Valdes-Sosa, P. a, Roebroeck, A., Daunizeau, J., Friston, K., 2011. Effective connectivity: influence, causality and biophysical modeling. Neuroimage 58, 339–61. https://doi.org/10.1016/j.neuroimage.2011.03.058

Welvaert, M., Rosseel, Y., 2013. On the definition of signal-to-noise ratio and contrast-to-noise ratio for fMRI data. PLoS One 8. https://doi.org/10.1371/journal.pone.0077089

Wen, X., Rangarajan, G., Ding, M., 2013. Is Granger Causality a Viable Technique for Analyzing fMRI Data? PLoS One 8. https://doi.org/10.1371/journal.pone.0067428

Winkler, I., Panknin, D., Bartz, D., Muller, K.R., Haufe, S., 2016. Validity of Time Reversal for Testing Granger Causality. IEEE Trans. Signal Process. 64, 2746–2760. https://doi.org/10.1109/TSP.2016.2531628

Zhang, J., Li, C., Jiang, T., 2016. New Insights into Signed Path Coefficient Granger Causality Analysis. Front. Neuroinform. 10. https://doi.org/10.3389/fninf.2016.00047

Zhou, D., Zhang, Y., Xiao, Y., Cai, D., 2014. Analysis of sampling artifacts on the Granger causality analysis for topology extraction of neuronal dynamics. Front. Comput. Neurosci. 8, 1–13. https://doi.org/10.3389/fncom.2014.00075

